# Muting, not fragmentation, of functional brain networks under general anesthesia

**DOI:** 10.1101/2020.07.08.188011

**Authors:** Corson N. Areshenkoff, Joseph Y. Nashed, R. Matthew Hutchison, Melina Hutchison, Ron Levy, Douglas J. Cook, Ravi S. Menon, Stefan Everling, Jason P. Gallivan

## Abstract

Changes in resting-state functional connectivity (rs-FC) under general anesthesia have been widely studied with the goal of identifying neural signatures of consciousness. This work has commonly revealed an apparent fragmentation of whole-brain network structure during unconsciousness, which has been interpreted as reflecting a break-down in connectivity and disruption in the brains ability to integrate information. Here we show, by studying rs-FC under varying depths of isoflurane-induced anesthesia in nonhuman primates, that this apparent fragmentation, rather than reflecting an actual change in network structure, can be simply explained as the result of a global reduction in FC. Specifically, by comparing the actual FC data to surrogate data sets that we derived to test competing hypotheses of how FC changes as a function of dose, we found that increases in whole-brain modularity and the number of network communities considered hallmarks of fragmentation are artifacts of constructing FC networks by thresholding based on correlation magnitude. Taken together, our findings suggest that deepening levels of unconsciousness are instead associated with the increasingly muted expression of functional networks, an observation that constrains current interpretations as to how anesthesia-induced FC changes map onto existing neurobiological theories of consciousness.

## I. INTRODUCTION

While much work has focused on how different anesthetics affect ion channels and receptor function at the cellular level (Anis et al. 1983, Franks 2006, Peduto et al. 1991), it remains poorly understood, by comparison, how anesthetics affect the coordinated activity of distributed whole-brain networks (Alkire et al. 2008, Brown et al. 2011). In recent years, resting-state functional MRI (rs-fMRI) has provided important glimpses into the large-scale, network-level effects of anesthesia. This approach, which measures covariance structure in spontaneous low-frequency oscillations in neural activity (Biswal et al. 1995), has repeatedly revealed an apparent fragmentation of functional brain network structure during various states of unconsciousness (Boly et al. 2012b, Hudetz and Mashour 2016, Hutchison et al. 2014). These findings are broadly consistent with work using electroencephalography and electrocorticography reporting a similar breakdown in long-range cortical communication under anesthesia (Lee et al. 2009) and during sleep (Ferrarelli et al. 2010).

Several authors (e.g. Boly et al. 2012b, Hudetz and Mashour 2016, Hutchison et al. 2014, Standage et al. 2019) have, implicitly or explicitly, interpreted these findings through the lens of various neurobiological theories of consciousness, which posit that conscious experience arises through the distributed processing of information throughout the neocortex. For example, the global neuronal workspace theory (Mashour et al. 2020) submits that information is made consciously accessible when it is broadcast widely throughout the cortex by a set of diffusely connected control regions in the prefrontal and parietal cortices. These regions in particular are frequently assigned a key role in the neural substrate of consciousness, and imaging research has revealed a reduction in frontal-parietal connectivity both during sleep (Spoormaker et al. 2012, Tagliazucchi et al. 2013) and under anesthesia (Boly et al. 2012a, Ku et al. 2011). By these, and other similar accounts (e.g. Tononi 2004), the fragmentation of network structure observed during sleep or anesthesia can be viewed as a causal signature of unconsciousness, resulting in a disruption of the brain’s ability to integrate information, or to broadcast it widely enough to create conscious awareness (Mashour 2013, Mashour and Hudetz 2018).

Network fragmentation is often concluded on the basis of changes in graph properties (e.g., increases in modularity or the number of communities) estimated from thresholded correlation matrices using a fixed, magnitude threshold. However, in this regime, global decreases – or “muting” – of functional connectivity may also produce the appearance of fragmentation by producing sparser, more disconnected networks (see Figure 1a). As an illustrative example, we contrast work which observed an increase in the modularity and number of communities of significance thresholded networks during sleep (Boly et al. 2012b), with work which found no such effect when using a relative threshold, chosen to obtain a fixed edge density (Uehara et al. 2014). It is diffcult to reconcile these results, or to establish that they represent truly distinct phenomena, using graph summary measures alone, as doing so requires a detailed analysis of the underlying network structure. This consideration is particularly pertinent in the case of anesthesia, which has been observed to result in overall decreases in correlation magnitude (Bettinardi et al. 2015, Lv et al. 2016). Further, while several authors have highlighted diminished long-range connectivity under anesthesia – notably between the frontal and parietal cortices (Hudetz and Mashour 2016) – other work has observed similar decreases in local connectivity (Monti et al. 2013). This suggests that anesthetic compounds may affect both short and long range cortical connections, and thus do not simply result in the disconnection of distant cortical regions, but disrupt coordinated neural activity at the local level (Hudetz et al. 2016, Vizuete et al. 2014).

**FIG. 1:**
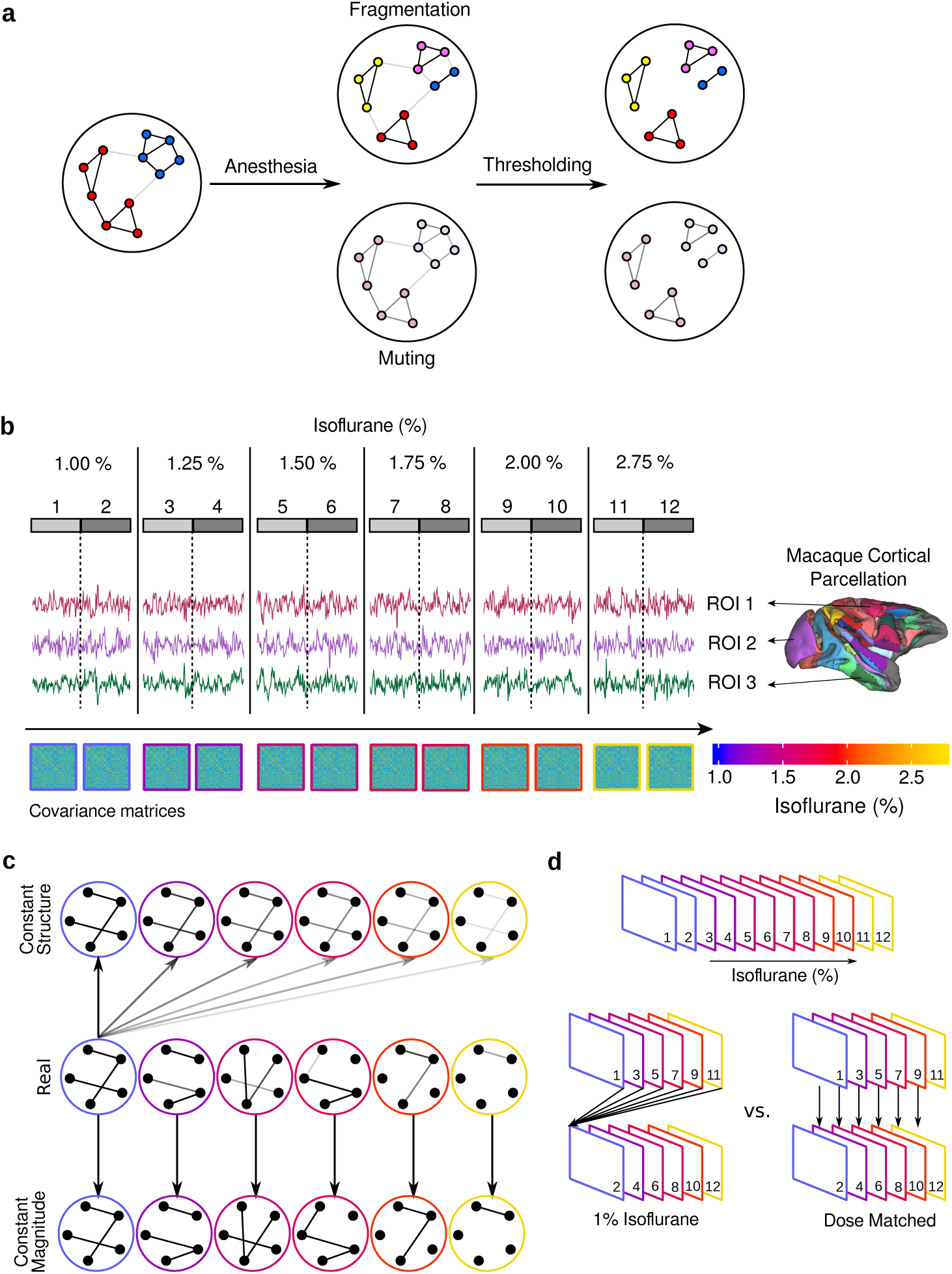
Competing hypotheses, methods and analytic approaches. **a)** Alternative accounts of the apparent network fragmentation observed during unconsciousness. The *fragmentation* account posits a splitting of conscious brain networks into smaller subnetworks during unconsciousness. An alternative *muting* account explains the apparent fragmentation by a global reduction in correlation magnitude, which results in sparser, more fragmented networks after applying a magnitude threshold. **b)** We collected twelve five-minute resting state scans – two at each of six concentrations of isoflurane. The cortex was parcellated using the LVE atlas (Lewis and Van Essen 2000), and covariance matrices were estimated for each scan. **c)** We studied dose related changes in network statistics both in the **real** data, and in surrogate datasets in which either the correlation structure or magnitude (the mean absolute value of the pairwise correlations) were held constant. In the former (**Constant Structure**), we simply scaled the correlation matrix associated with the lowest dose (1% isoflurane) to match the observed mean magnitude for each scan. In the second (**Constant Magnitude**), we scaled all correlation matrices to have mean magnitude equal to the lowest dose. **d)** To quantify the degree to which the data supported either a constant vs. dose dependent covariance structure, we split each subjects data in two datasets, each comprising one of the two scans at each dose. We then compared the correlation matrices in one half both to the lowest dose (**1% isoflurane**) or to the corresponding dose (**Dose matched**) in the other.

For studies seeking to identify the neural correlates of unconsciousness using functional connectivity, these facts suggest an important distinction – between alterations in network structure on one hand, versus overall changes in correlation magnitude on the other. These different effects can be difficult to disentangle, as evidenced by findings of both (1) an increase in the modularity and number of communities during sleep, and (2) that, despite these network changes, overall network structure remained relatively preserved (Boly et al. 2012b). Given this general ambiguity, as well as the ambiguity as to how such results may support theories of consciousness (Tononi 2004), the goals of the present study were two-fold: First, to ascertain to what extent network fragmentation under anesthesia-induced unconsciousness is attributable to an overall decrease in connectivity strength; and second, to characterize the structure of whole-brain functional connectivity across depths of unconsciousness. To unpack these relationships, we examined changes in rs-fMRI brain network structure in non-human primates across six increasing levels of anesthesia. This allowed us to test for fine-graded changes in network strength and structure across sedation levels, while also assessing critical components of neurobiological theories related to consciousness.

## II. RESULTS

### A. Reduction in correlation magnitude explains apparent network fragmentation

Figure 2a displays summary statistics for the correlation matrices estimated from each scan. Consistent with previous work (Xie et al. 2019), increasing dose was associated with an overall decrease in correlation magnitude, with no clear change in the ratio of positive to negative correlations (as in Bettinardi et al. 2015). We also observed a decrease in the spectral radius (the magnitude of the largest eigenvalue), suggesting an overall loss of low dimensional structure. This reduction in correlation did not appear to be driven by regions in any single network, but was present both in primary sensory and somatomotor regions, as well as in the frontal and parietal cortices, and connections between them (Figures 2b-c). Although bilateral homologs displayed relatively high functional connectivity compared to non-homologous regions at low doses, this connectivity was likewise seen to sharply decrease with increasing dose (Figure 2d). Figures 2e-f displays the decrease in correlation magnitude at the level of individual ROIs, suggesting that all regions show a trend towards decreasing mean connectivity with increasing dose. Together, these effects indicate a global reduction in functional connectivity magnitude as isoflurane dose increases.

**FIG. 2:**
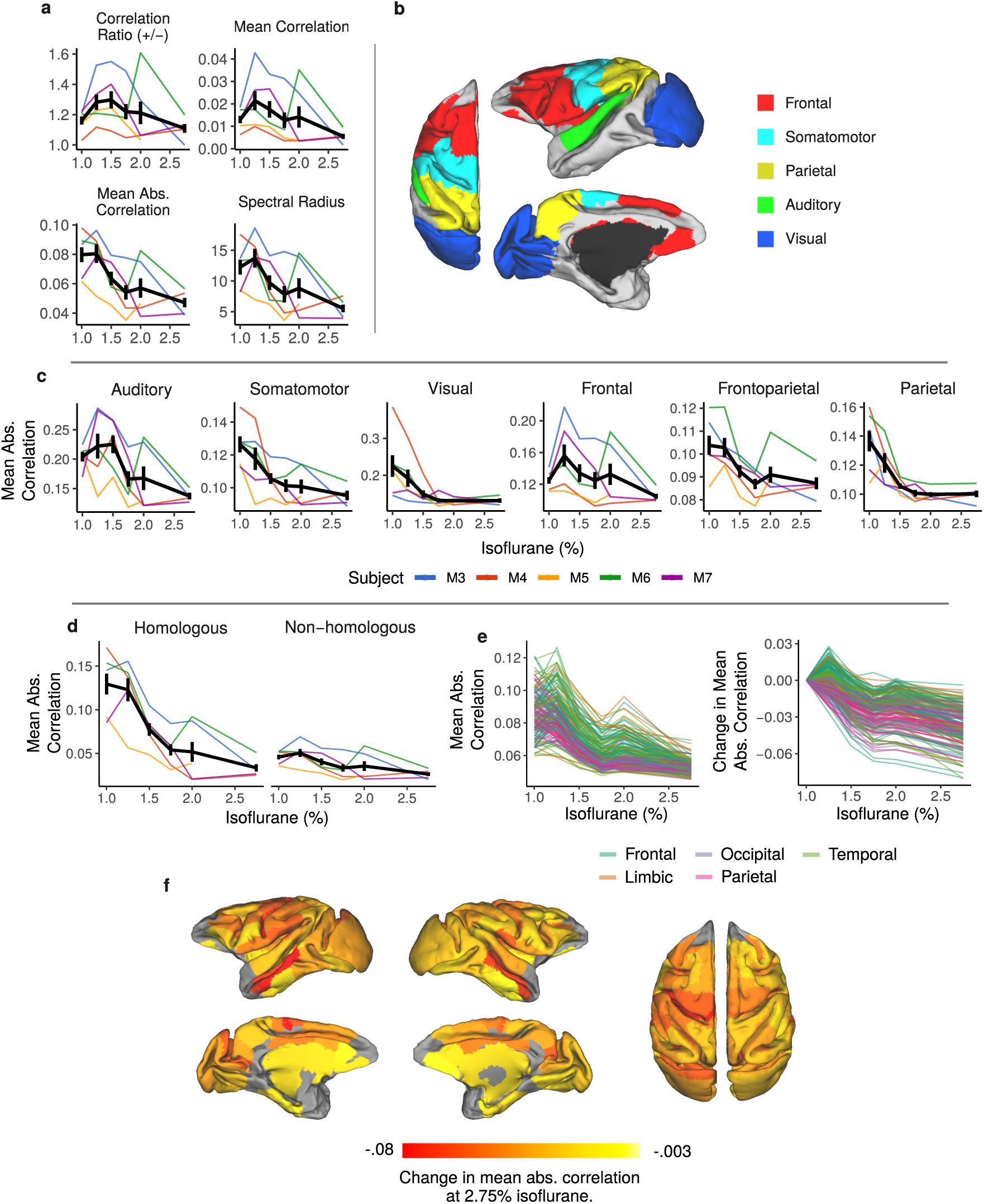
Correlation magnitude decreases both globally and locally as a function of increasing anesthetic dose. **a)** Descriptive statistics for correlation matrices. In all figures, colored lines denote means for each each subject. Solid black line denotes the mean across all subjects. Error bars are one standard error. **b)** Parcelation of the cortex into primary sensory regions (somatomotor, auditory, and visual), and frontal and parietal cortices. **c)** Mean absolute correlation within each parcel, as well as between the frontal and parietal cortices (frontoparietal). **d)** Mean absolute correlation between bilateral homologues and non-homologous ROIs. **e)** Average muting of functional connectivity per ROI. Figures display the mean absolute correlation, as well as the change in mean absolute correlation relative to the lowest dose (1% isoflurane). **f)** Surface map of the change in mean absolute correlation at 2.75% isoflurane relative to the lowest dose.

Next, to determine the extent to which network changes across dose can be attributed to actual changes in network structure, versus an overall decrease in correlation magnitude, we computed graph summary statistics from the thresholded and binarized correlation matrices, as well as from two surrogate datasets in which either the average correlation structure or the correlation magnitude was held constant across dose. These surrogate datasets were important, as they provided a critical basis for interpreting effects in the real data; if the appearance of network fragmentation is due primarily to an overall decrease in correlation magnitude, then the same pattern of results should be observed when the exact same correlation structure is held constant across doses, and only the magnitude is allowed to vary. Conversely, fragmentation should be abolished when correlation matrices are scaled to have the same average magnitude. We created the **Constant Structure** surrogate dataset by replacing each correlation matrix (from 1.00% – 2.75% isoflurane) with a copy of the average of the subjects two correlation matrices at the lowest dose (1.00%), scaled to have matching correlation magnitude to the real data (defined as the mean absolute value over all pairwise correlations). By contrast, we created the **Constant Magnitude** surrogate dataset by scaling each correlation matrix to have the same average correlation magnitude as in the corresponding subjects lowest dose condition (see Figure 1c). We then thresholded and binarized each correlation matrix using an uncorrected, one-tailed t-test with a significance threshold of *α* = .05. For further comparison, we also binarized the observed correlation matrices using a relative threshold chosen to produce an constant edge density equal to the mean edge density at the lowest dose (**Density Threshold**). If increasing dose is characterized primarily by a overall decrease in correlation strengths, rather than a change in network structure, then graph properties of density thresholded networks should be relatively preserved across dose.

Broadly consistent with previous work (Boly et al. 2012b, Hutchison et al. 2014) – and the interpretation that brain networks become increasingly fragmented and/or disconnected at increased levels of sedation – we observed an increase in sparsity, the number of communities, modularity, and small-worldness with increasing dose, along with a decrease in network efficiency (Figure 3a). Notably, however, we observed a nearly identical pattern of results in our surrogate dataset in which the correlation structure was held constant across dose (Constant Structure), and only the correlation magnitude was varied. Furthermore, these effects were completely abolished in our surrogate dataset in which the correlations were scaled to have common magnitude across dose, or when the networks were thresholded to achieve a fixed edge density. These results, when taken together, suggest that the observed effect of increasing dose on network statistics (i.e., fragmentation) is better explained as a reduction in overall correlation magnitude, or the muting of constant network structure.

**FIG. 3:**
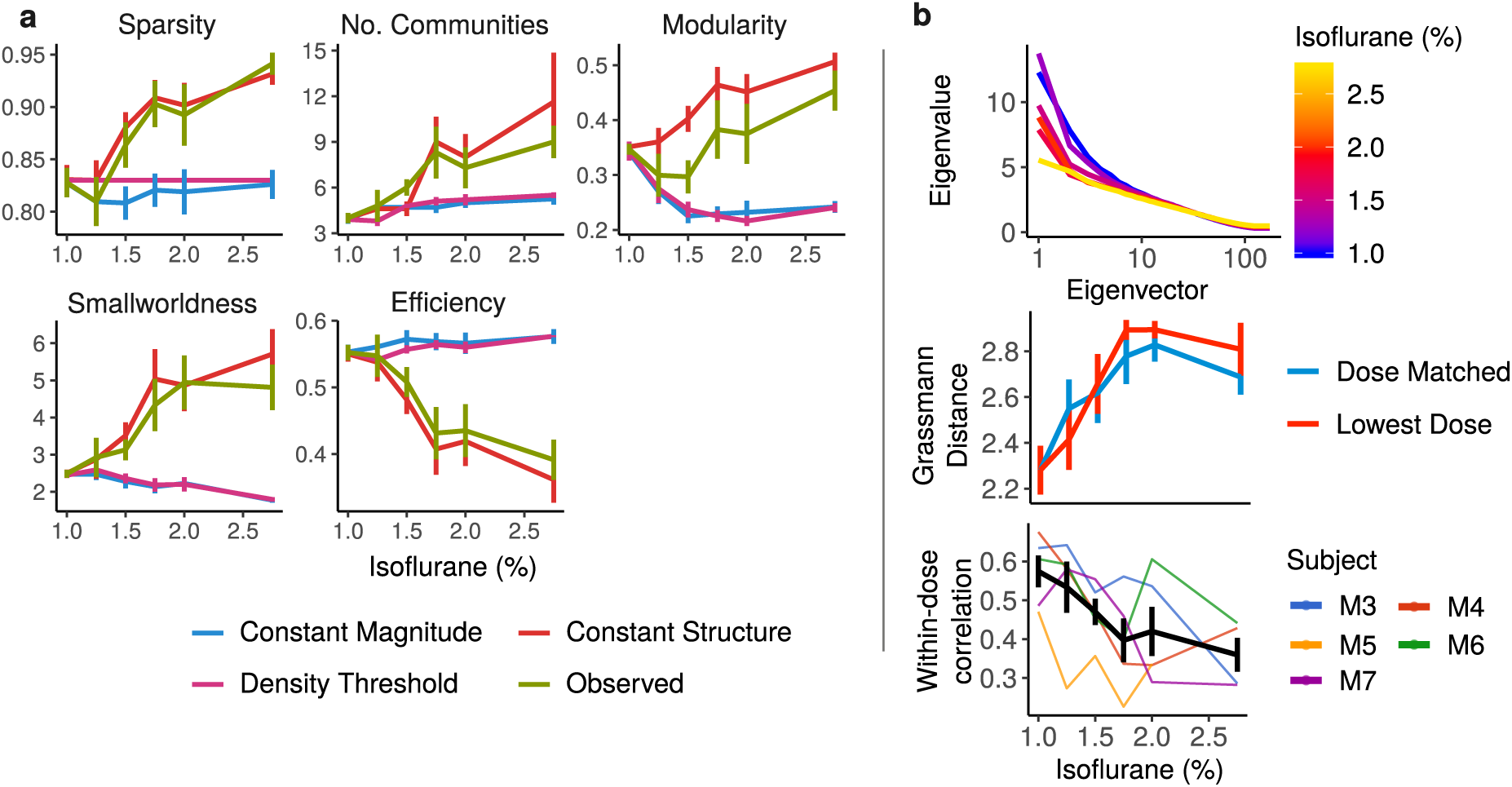
The appearance of network fragmentation is an artifact of an increasingly muted network structure. **a)** Network summary statistics. Values denote means across all subjects, while error bars denote one standard error. **Observed** denotes the actual, observed correlation matrices. For **Constant Magnitude**, correlations at each dose level were scaled to match the average magnitude of the correlations at the lowest dose (1% isoflurane). For **Constant Structure**, the correlation matrix for the lowest dose was replicated across all scans, and scaled to match the observed average magnitude. Networks were constructed by thresholding and binarizing correlation matrices by significance using a one-tailed t-test with *α* = .05. For comparison, we also thresholded the observed correlation matrices using a relative threshold to achieve a fixed edge density of .3 (**Density threshold**). Communities were estimated using the Louvain clustering algorithm. Note that the effect of isoflurane on the observed networks is consistent with the scaling of a constant correlation structure. **b)** Mean spectrum of the correlation matrices at each dose (top). The x-axis is shown on a log scale to better display the effects of dose on the leading eigenvalues. After splitting each subjects data into two halves – comprising the first and second scans at each dose, respectively – we compared the correlation matrices in first half either to the corresponding dose (**Dose Matched**), or to the lowest dose (**1% Isoflurane**) in the other. Correlation matrices were compared using the distance between the subspaces spanned by the leading five eigenvectors (middle). The bottom figure displays the correlation between the vectorized correlation matrices from the two scans at each dose, suggesting that the reliability of functional connectivity estimates decreases with increases depth of anesthesia.

### B. Dose effects are well explained by a constant network structure

If the observed dose effects can be explained by the muting of a constant network structure, then the correlation structure at all dose levels should be well approximated by the structure at the lowest dose (1% isoflurane). We directly tested this by splitting each subject’s data into two halves, comprising the first and second scans at each dose, respectively. We then compared each whole-brain correlation matrix in the first set either to the corresponding dose in the second, or to the lowest dose, using the distance between the subspaces spanned by the leading eigenvectors (the geodesic distance on the Grassmann manifold; Edelman et al. 1998). We chose the subspace spanned by the leading five eigenvectors, as this was the range of the spectrum most strongly affected by dose (Figure 3b, top).

Critically, we found a near identical pattern of results in the lowest dose and in the dose matched comparisons (Figure 3b, middle), indicating that the dominant patterns of network structure at higher dosages are well approximated by the structure already present at the lowest dose.

To visualize this structure, we used common principal component analysis (CPCA; Flury 1984, Trendafilov 2010) to derive a set of components summarizing the correlation structure across dosages. Prior to applying CPCA, subject correlation matrices were centered to remove subject differences in functional connectivity. This decision was motivated by previous findings (Gratton et al. 2018, Xu et al. 2019) that variability in functional connectivity in Humans and non-human primates is dominated by stable, subject level effects, which may mask the relatively small differences induced by task manipulations. Consistent with these findings, we found strong clustering of the correlation matrices at the subject level (Figure 4a, left) in the uncentered data, which we were able to remove through our centring approach (Figure 4a, right).

**FIG. 4:**
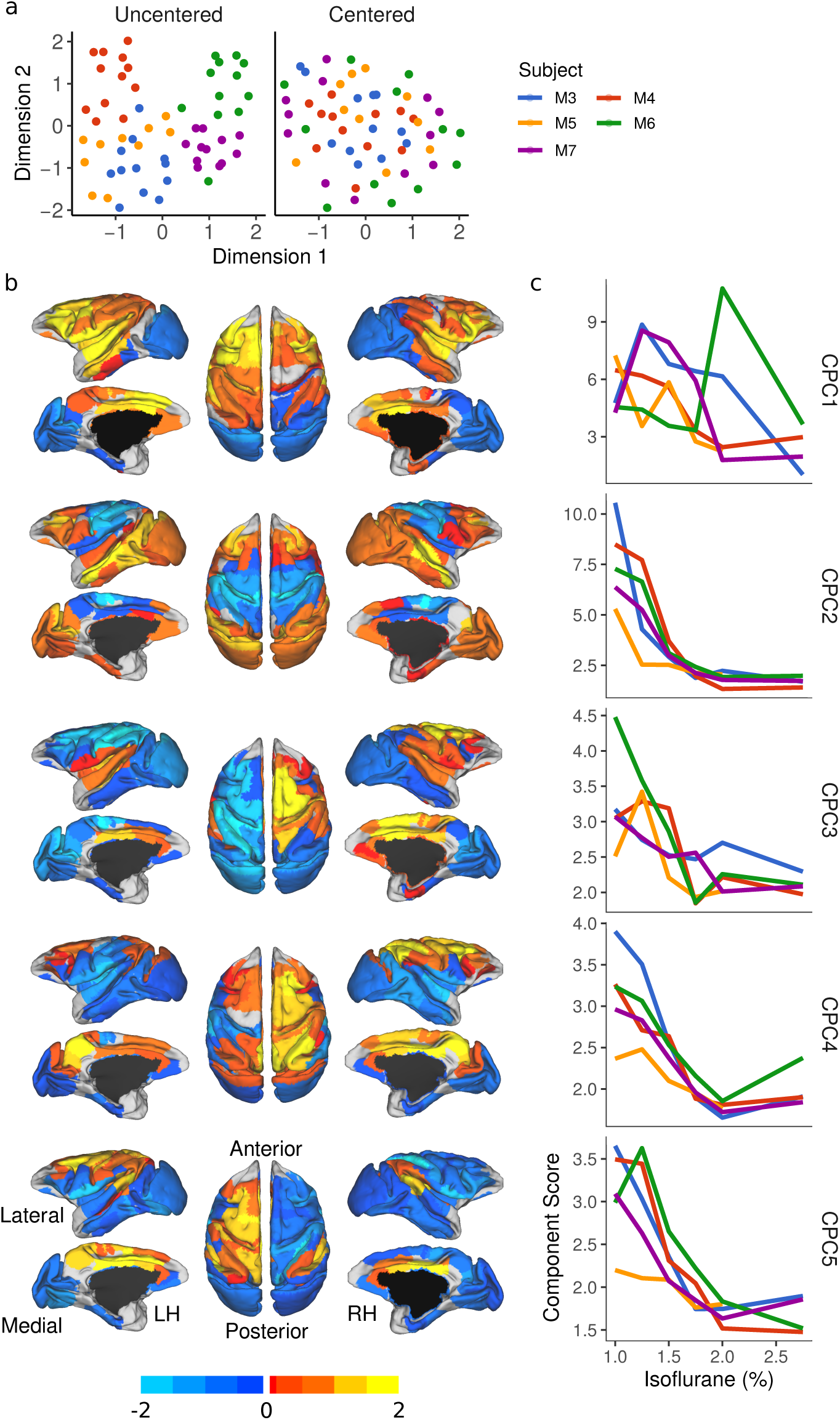
Latent structure of brain networks is present across all dose levels, but becomes increasingly muted. Common principal component analysis (CPCA) of subjects’ centered covariance matrices. The number of displayed components was selected on the examination of the spectrum of the observed correlation matrices (see Figure 3b). Prior to CPCA, subject covariance matrices were centered to remove static subject differences in functional connectivity. **a)** Two-dimensional embedding of subject covariance matrices by uniform manifold approximation (UMAP; McInnes et al. 2018) using a distance matrix constructed by the pairwise geodesic distances. Note the subject level clustering in the uncentered data. **b)**. Spatial maps for the top five components. **c)** Component scores for each subject and each scan.

Figure 4b-c shows the spatial maps of the components extracted by applying CPCA to the centered correlation matrices, as well as the contributions of these components at each dose. The first of these (CPC1) constitutes a gradient separating visual and medial ventrotemporal areas from parietal, frontal and superior temporal regions. CPC2 constitutes a gradient separating somatomotor areas from the rest of the cortex. CPC3 constitutes a gradient separating right parieto-frontal cortex and bilateral superior temporal cortex from the rest of cortex; CPC5 is a near-mirror image of this same gradient; finally CPC4 constitutes a gradient separating bilateral frontoparietal cortex from the rest of the cortex. The top components are consistent with some of the gradients reported by others in both non-human primates (Margulies et al. 2016, Yacoub et al. 2020), and in humans (Hong et al. 2020). Together, these findings support the notion that a constant network structure is present across all dose levels, and that it is only the expression of this constant network structure that changes across dose. With respect to the latter, indeed we find that these components decreased almost monotonically with increasing dose, asymptoting at approximately 2% isoflurane.

## III. DISCUSSION

We analyzed static resting state functional connectivity (rs-FC) under increasing depths of anesthesia in order to characterize dose-related changes in cortical whole-brain network structure. Increasing dose was associated with an increase in network modularity and the number of communities, as well as a decrease in network efficiency, all of which is consistent with the apparent fragmentation discussed in previous literature (Hudetz and Mashour 2016). However, comparisons with surrogate datasets in which either the correlation structure or magnitude were held constant revealed that these effects, rather than reflecting a qualitative change in network structure (i.e., network fragmentation), could be fully explained as an overall reduction in correlation magnitude, which we refer to as muting. Further supporting the idea that network structure was unchanged across dose levels, we showed that the principal components of the functional connectivity matrices at each increasing dose level (from 1.00% – 2.75%) were just as similar to the components at the lowest dose (1.00% isoflurane) as they were to the components in another, dose-matched scan. This suggests that there is no explanatory benefit to assuming a change in correlation structure across dose levels, as this structure can be just as well approximated assuming a fixed structure, already present at the lowest dose. Finally, we used common principal component analysis to derive a set of components that summarize the correlation structure across dose, and show that the expression of these components decreases in a near monotonic fashion as dose increases. Taken together, these findings suggest that deepening levels of unconsciousness are associated with the increasingly muted expression of a constant functional network structure, rather than a break-down or fragmentation of this structure. We discuss both the methodological and theoretical implications of these findings below.

Several neurobiological theories of consciousness center on the brains capacity for information exchange across multiple distributed regions, with unconsciousness being a result of the disruption of this information transmission (Mashour et al. 2020). For instance, the global neuronal workspace model posits that information, initially encoded by a specialized processing unit, is made consciously accessible when it is broadcast widely throughout the cortex, to the many other specialized processors in the brain (Mashour et al. 2020). By association, conditions that impact this global broadcasting of information, from sleep and anesthesia to brain damage, can result in varying levels of unconsciousness (Mashour and Hudetz 2018). Graph theory measures, such as modularity and network efficiency, have routinely been utilized as indirect estimates of the capacity for information transfer across brain networks, and have often been used to draw inferences concerning changes in network structure during unconsciousness. For instance, a reduction in network efficiency has been observed in both local and global brain networks during anesthetic- and sedative-induced unconsciousness, and has been interpreted as reflecting a dysfunction in communication across the cortex (Hashmi et al. 2017, Monti et al. 2013).

Consistent with this interpretation, decreases in local and global efficiency have also been observed in patients with consciousness disorders, such as unresponsive wakefulness syndrome and the minimally conscious state, and such decreases have been shown to correlate with reductions in patient awareness (Chennu et al. 2014). Likewise, increases in whole-brain modularity and the number of network communities have been observed during both non-rapid eye movement sleep (Boly et al. 2012b) and isoflurane-induced anesthesia (Hutchison et al. 2014, Standage et al. 2019), which has been interpreted as reflecting a literal fragmentation of brain networks into smaller, more isolated processing units. These interpretations are understandably compelling, as graph measures are thought to quantify key aspects of information transmission in the brain, and many of them, such as modularity and efficiency, neatly map onto existing frameworks and hypotheses concerning theories of consciousness (and disruptions thereof).

In the current study, we have demonstrated that changes in graph metrics; such as whole-brain modularity, network efficiency, path length, and number of communities, can all be simply explained by an overall reduction in functional connectivity, which we refer to as muting. This contrasts markedly with the way that changes in graph metrics have been commonly interpreted, which is that they reflect some qualitative change in underlying brain network structure or organization. Distinguishing between these two phenomena (muting vs. fragmentation) requires a detailed analysis of the change in network structure across levels of anesthesia. As we have shown here, graph summary statistics are highly sensitive both to the threshold techniques used to construct the networks, and to overall changes in correlation strength. This makes it difficult to draw firm conclusions about changes in network structure from graph statistics alone. As demonstrated with our surrogate data, the community structure of a thresholded correlation network can be altered in a manner consistent with fragmentation, even when the overall pattern of correlations is held constant. This is consistent with what we observe in the real data: using CPCA, we demonstrate that, despite a supposed network fractionation (i.e., increased modularity and number of communities), the exact same network structure is present across all dose levels, with only the expression of this structure being weakened (or ‘muted’) at higher levels of sedation. We also note although we are wary of drawing conclusions from a single observation that subject M5, which experienced an adverse reaction to the anesthetic at the highest dose, showed a relatively flat response to isoflurane, displaying functional connectivity consistent with higher doses even at the lowest dose (Figure 4). This may suggest that hypersensitivity to isoflurane, or the potential for adverse reactions, are detectable in functional connectivity even at low doses.

One of the main observations to emerge in the anesthesia literature, and which has been frequently used to bolster neurobiological theories of consciousness, is that disruptions in consciousness are associated with a break-down in long-range frontal-parietal connectivity. We observed similar reductions in frontal-parietal connectivity under increasing depths of anesthesia; notably, however, this effect was not unique to long-range connectivity, as we also observed comparable decreases within both the frontal and parietal cortices, as well as the primary sensory (visual, auditory) and somatomotor (primary sensory and motor) areas. Taken together, these findings bolster our interpretation that decreases in long-range functional connectivity do not reflect a literal fragmentation of these cortices into distinct networks, as much as it is a consequence of a brain-wide, global reduction in correlation magnitude. This is consistent with previous fundings that propofol anesthesia produces disruptions in local as well as long-range connectivity (Monti et al. 2013). While this seemingly departs from the view that long-range frontal-parietal networks play a unique role in conscious experience per se, it nevertheless comports with the broader view, shared by several theories, that unconsciousness stems from a more global disruption of information processing throughout the brain. It is also consistent with views which assign frontal-parietal regions a priviledged position due to their role as hub regions facilitating the broadcasting of information widely across the cortex (Mashour et al. 2020), as this broadcasting may be impaired not only by selective disruption of frontal-parietal regions, but also by more global impairments in connectivity.

Our finding that functional connectivity is decreased even in sensory regions is consistent with work showing that anesthetic compounds interfere with neural synchronization even in these regions. For example, local field potentials in the visual cortex form spontaneous, spatially localized patterns of synchronized activity which grow more variable and entropic under anesthesia (Hudetz et al. 2016), and local firing patterns become decorrelated (Vizuete et al. 2014). These local disruptions in coordinated neural activity may result in noise manifesting as a reduction in functional connectivity. Thus, the results we have observed may reflect not simply a reduction in communication within and between brain networks, but the fading of an existing network structure into a background of neural noise. This account would also be consistent with our observation that increasing depth of anesthesia was associated with a flattening of the spectrum in the estimated correlation matrices, but with relatively little change in the dominant eigenstructure (Figure 3, top and middle). As well as our observation that the reliability of functional connectivity estimates decreased with increasing dose (Figure 3, bottom).

Our findings should be interpreted in light of a few methodological considerations. First, our study did not collect neuroimaging data from monkeys during the awake state, limiting our discussion and interpretations to the network structure present across deepening levels of anesthesia (and presumably unconsciousness). However, Bettinardi et al. (2015) note a gradual emergence of coordinated brain activity during the transition from deep sedation to wakefulness, suggesting that our protocol nonetheless samples a portion of a continuous trajectory spanning waking and unconsciousness. Second, we induced unconsciousness through isoflurane (Hutchison et al. 2014, Shmuel and Leopold 2008, Vincent et al. 2007), which is one of only several different anesthetics that could have been used. Isoflurane is a potent vasodilator (Iida et al. 1998), and has been shown to have effects on cerebral blood flow and blood volume (Li et al. 2013, Masamoto et al. 2006). As such, it is possible that its hemodynamic properties could obscure potential neural changes that occur at higher dose levels. However, studies combining fMRI with electrical recordings, such as EEG-fMRI (Barttfeld et al. 2015, He et al. 2008, Liu et al. 2013, Ranft et al. 2016, Vincent et al. 2007), have demonstrated a close coupling between changes in neural activity and corresponding changes in functional connectivity. Moreover, our finding of reduced functional connectivity at both the small and large-scale is consistent with similar observations under propofol (Monti et al. 2013), and studies examining network dynamics under anesthesia have observed similar effects under several different types of anesthesia, from propofol and sevoflurane to ketamine (Barttfeld et al. 2015, Hutchison et al. 2014, Uhrig et al. 2018). Thus, we think it unlikely that our findings are entirely explainable by the specific mechanisms of action of the anesthetic used here.

Theories relating consciousness to the information processing capacity of the brain must ultimately generate concrete predictions describing the patterns of brain activity supporting conscious awareness. Functional neuroimaging plays a central role in the testing of these predictions, as these techniques allow for the precise characterization of brain activity across various states of consciousness and unconsciousness. Although fMRI affords the opportunity to study the largescale, structural features of whole-brain networks, the complexity of this data necessitates careful analysis. As we have shown, graph summary measures lack the resolution to fully characterize whole-brain network structure, or changes in this structure across depths of unconsciousness. Further, they are highly sensitive to the methods used to construct the networks, making it difficult to determine what precise properties of the brain are being quantified by these measures. Our results suggests that a global muting of functional connectivity is a significant feature of isoflurane induced anesthesia, and that this fact is sufficient to explain previously reported results which are commonly attributed to structural changes in whole-brain networks. They also highlight the need to carefully disentangle putative changes in network structure from effects induced by overall changes in correlation magnitude, or by the method of network construction; something that is likely to be difficult through the use of graph summary measures alone. We have not addressed the causal role of this muting, or whether it may simply mask structural network features underlying consciousness; although recent work (Pal et al. 2020) suggests that anesthesia induced unconsciousness can be dissociated from the effects of anesthesia on the cortex, and so the relationship between consciousness and cortical information processing may be more complex than can be decoded from BOLD signal correlations alone.

## IV. METHODS

### A. Data collection

We reanalyzed data from five Macaque primates (M. Fascicularis; 4 female; Mean age 7.8 yrs) collected as part of the experiment reported in Hutchison et al. (2014). All surgical and experimental procedures were carried out in accordance with the Canadian Council of Animal Care policy on the use of laboratory animals and approved by the Animal Use Subcommittee of the University of Western Ontario Council on Animal Care. As data acquisition is thoroughly described by the original authors, we present a more condensed description here. Note however, that our fMRI preprocessing pipeline contains slight differences.

Prior to image acquisition, subjects were injected intramuscularly with atropine (0.4 mg/kg), ipratropium (0.025 mg/kg), and ketamine hydrochloride (7.5 mg/kg), followed by intravenous administration of 3 ml propofol (10 mg/ml) via the saphenous vein. Subjects were then intubated and switched to 1.5% isoflurane mixed with medical air. Each subject was then placed in a custom-built chair and inserted into the magnet bore, at which time the isoflurane level was lowered to 1.00%. Prior to image localization, shimming, and echo-planar imaging (EPI), at least 30 min was allowed for the isoflurane level and global hemodynamics to stabilize at this concentration. We then acquired 2 functional EPI scans at each of six increasing isoflurane levels: 1.00, 1.25, 1.50, 1.75, 2.00, and 2.75% (0.78, 0.98, 1.17, 1.37, 1.56, and 2.15 minimum alveolar concentration, respectively). We interleaved a 10 min period between each isoflurane level increase to allow for the concentration to stabilize, during which no fMRI data were collected. Throughout the duration of scanning, the monkeys spontaneously ventilated and we monitored physiological parameters (temperature, oxygen saturation, heart rate, respiration, and end-tidal CO2) to ensure that values were within normal limits. The acquisitions of two anatomical images occurred during the stabilization periods between isoflurane levels.

The monkeys were scanned on an actively shielded 7-Tesla 68-cm horizontal bore scanner with a DirectDrive console (Agilent, Santa Clara, California) with a Siemens AC84 gradient subsystem (Erlangen, Germany). We used a custom in-house conformal five-channel transceive primate-head Radio Frequency (RF) coil. Each functional run consisted of 150 continuous EPI functional volumes (repetition time [TR] = 2000 ms; echo time [TE] = 16 ms; flip angle = 700; slices = 36; matrix = 96 × 96; Field of view [FOV] = 96 × 96 mm2; acquisition voxel size = 1 × 1 × 1 mm3), acquired with GRAPPA = 2. A high-resolution gradient-echo T2 anatomical image was acquired along the same orientation as the functional images (TR = 1100 ms, TE = 8 ms, matrix = 256 × 256, FOV = 96 × 96 mm2, acquisition voxel size = 375 × 375 × 1,000 mm3). We also acquired a T1-weighted anatomical image (TE = 2.5 ms, TR = 2300 ms, FOV = 96 × 96 mm2, acquisition voxel size = 750 × 750 × 750 mm3).

### B. fMRI preprocessing

Functional image preprocessing was implemented using Nipype (Neuroimaging in Python: Pipelines and Interfaces; http://nipy.org/nipype). The functional images underwent de-spiking, motion correction (six-parameter affine transformation), and slice-time correction, before brain extraction, and highpass temporal filtering (0.01 Hz; Hallquist et al. 2013, Power et al. 2014). Functional data was co-registered to its respective T1 anatomical (six degrees of freedom rigid transformation), and then linear (12 degrees of freedom linear affine transformation) and nonlinear transformed to the F99-template (Van Essen 2002) and parcellated into 174 regions of interest (ROI) using the LVE atlas (Lewis and Van Essen 2000). All further analyses were done in R (version 3.6.1; R Core Team 2019).

Nuisance regression was performed using the six motion parameters, their derivatives, their squares, as well as mean white matter and CSF signals, for a total of 26 regressors. We elected to use the unwhitened residuals rather than performing prewhitening, as the overall residual auto-correlation was minimal (mean Durbin-Watson statistic in the range [1.6-2] for all subjects and all scans), and did not appear to be related to dose.

### C. Functional connectivity estimation and comparison

For each scan, we estimated a covariance matrix from the standardized residuals using the shrinkage estimator proposed by Ledoit and Wolf (2004), which has been shown to perform better than the sample covariance matrix in high-dimensional, small-sample settings.

Networks were constructed from the real and surrogate datasets by thresholding the entries of each correlation matrix using a one-tailed t-test with a threshold of *α* = .05. All graph metrics (Figure 3a) save smallworldness were computed from the binarized correlation matrices using the R package igraph (Csardi and Nepusz 2006), and community detection was performed using the Louvain clustering algorithm (Blondel et al. 2008). Smallworldness was computed using the R package brainGraph (Watson 2019).

Structural similarity between correlation matrices was quantified using the distance between the subspaces spanned by the leading five eigenvectors (Figure 3b). These five eigenvectors constitute a basis for a five-dimensional subspace of the full space of 174 ROIs. The set of all such subspaces forms a manifold – called the Grassmann manifold – on which several natural distance measures can be defined (Edelman et al. 1998). We define the distance between two such subspaces to be the arc length of the geodesic between them, equivalent to the magnitude of the vector of principle angles between the subspaces. Specifically, let **V**_1_ and **V**_2_ be matrices whose columns are the top *k* eigenvectors of the correlation matrices **S**_1_ and **S**_2_, respectively. Then the principal angles between the subspaces spanned by **V**_1_ and **V**_2_ are given by *θ*_*i*_ = cos^−1^ *σ*_*i*_, where *σ* = {*σ*_1_, …, *σ*_*k*_} are the singular values of 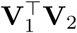. The distance between **V**_1_ and **V**_2_ is then

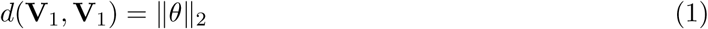

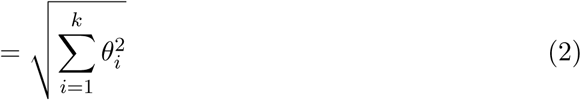

### D. Centering

To center each subject’s correlation matrices, we took the approach advocated by Zhao et al. (2018), which leverages the natural geometry of the space of covariance matrices. We have implemented many of the computations required to replicate the analysis in an R package **spdm** (**s**ymmetric **p**ositive-**d**efinite **m**atrix), which is freely available from a Git repository at https://gitlab.com/fmriToolkit/spdm.

The procedure is as follows. For each subject *i*, we computed a geometric mean covariance matrix 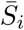 using the fixed-point algorithm described by Congedo et al. (2017), as well as a grand mean 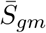 over all subjects and all scans. We then projected each covariance matrix *S*_*ij*_ onto the tangent space at the corresponding subject mean 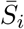 to obtain a tangent vector

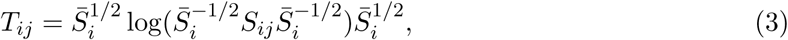

where log denotes the matrix logarithm. We then transported each tangent vector to the grand mean 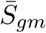 using the transport proposed by Zhao et al. (2018), obtaining a centered tangent vector

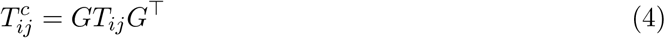

where 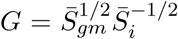. Finally, we projected each centered tangent vector back onto the space of covariance matrices, to obtain the centered covariance matrix

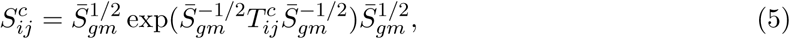

where exp denotes the matrix exponential.

To gain some intuition for this procedure, note that centering a sample of real numbers can be viewed as the process of computing a sample mean, and then applying a translation which takes the sample mean to zero. The resulting centered values can then be viewed as vectors describing the deviations of the observed values from the mean. In the same way, the tangent vectors computed in Eq. 3 can be viewed as the deviations of each correlation matrix from the corresponding subject mean. The transport in Eq. 4 translates these deviations to the grand mean, thus aligning them to a common baseline. A naive approximation to this procedure would involve, for each subject, simply subtracting the mean covariance matrix from the covariance matrix in each scan. This approach has several drawbacks; most importantly, the fact that the difference between two covariance matrices is not necessarily a covariance matrix, and so does not have an obvious interpretation. The procedure we have described above can be viewed as an adaptation of this approach, which respects the geometric structure of the space of covariance matrices.

To visualize the effect of centering (Figure 4a), we derived a two-dimensional embedding of the set of covariance matrices before and after centering using uniform manifold approximation (UMAP; McInnes et al. 2018). This procedure was applied to the distance matrices computed from the pairwise distances between covariance matrices (Smith 2005), defined as

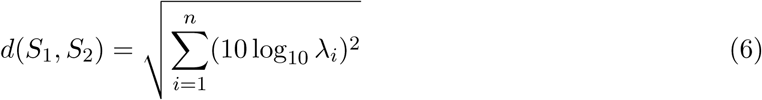

where (*λ*_1_, …, *λ*_*n*_) are the generalized eigenvalues of *S*_1_ and *S*_2_.

### E. Common component analysis

After centering, we sought an interpretable, low-dimensional summary of the observed covariance matrices in order to characterize changes in functional connectivity across depths of anesthesia. For a single covariance matrix, this could be accomplished by principal component analysis (PCA), where the eigenvectors of the covariance matrix are used as a basis for a low dimensional subspace capturing the dominant patterns of variability in the BOLD signal observed during a single scan. As we had observations for multiple scans and multiple subjects, we considered two approaches for simultaneously decomposing the full set of covariance matrices.

The first is common principal component analysis (CPCA; Flury 1984, Trendafilov 2010). The CPCA model attempts to simultaneously diagonalize multiple covariance matrices, and so (informally) assumes that the covariance matrices have identical factor structure, though they may differ in the degree to which they express those factors. As this is a restrictive assumption, we also considered a second model – the common component analysis (CCA) proposed by Wang et al. (2011) – which relaxes the assumption that the set of covariance matrices may be simultaneously diagonalized at the expense of some interpretability. The CPCA model was fit using the R package cpca (Ziyatdinov et al. 2014), while the CCA model was fit using custom R code implementing the iterative algorithm proposed by Wang et al. (2011). As both models returned highly similar results, we present only the results of CPCA.

